# Close encounters of three kinds: impacts of leg, wing, and body collisions on flight performance in carpenter bees

**DOI:** 10.1101/2022.10.21.513269

**Authors:** Nicholas P. Burnett, Stacey A. Combes

## Abstract

Flying insects often forage among cluttered vegetation that forms a series of obstacles in their flight path. Recent studies have focused on behaviors needed to navigate clutter while avoiding all physical contact, and as a result, we know little about flight behaviors that do involve encounters with obstacles. Here, we challenged carpenter bees (*Xylocopa varipuncta*) to fly through narrow gaps in an obstacle course to determine the kinds of obstacle encounters they experience, as well as the consequences for flight performance. We observed three kinds of encounters: leg, body, and wing collisions. Wing collisions occurred most frequently (in about 40% of flights, up to 25 times per flight) but these had little effect on flight speed or body orientation. In contrast, body and leg collisions, which each occurred in about 20% of flights (1-2 times per flight), resulted in decreased flight speeds and increased rates of body rotation (yaw). Wing and body collisions, but not leg collisions, were more likely to occur in wind versus still air. Thus, physical encounters with obstacles may be a frequent occurrence for insects flying in some environments, and the immediate effects of these encounters on flight performance depends on the body part involved.

## Introduction

Flying animals frequently interact with cluttered vegetation in their habitats. Many birds, for instance, nest and perch in trees and pursue prey through dense foliage [1,2], and many insects forage for nectar and pollen among dense patches of flowers [3,4]. In each case, animals navigate around a series of vegetative structures that functionally serve as obstacles and constrain navigable paths [5,6]. Traversing obstacles while in flight requires coordinated detection of obstacles (e.g., visually) and rapid alteration of the flight path, for example by decelerating, accelerating, or changing body orientation [7–11]. However, most studies of obstacle traversal in flight focus on behaviors required to completely avoid obstacles, with little consideration of what happens when animals do make contact with obstacles (e.g., collisions). This contrasts with studies of terrestrial locomotion that consider obstacle encounters as an integral part of traversing terrestrial landscapes [12–14]. Thus, we know relatively little about the effects of physical encounters with obstacles on the performance of flying animals.

Encounters with obstacles can alter locomotory performance at the time of the collision and also lead to performance-altering injuries that may be immediate or cumulative. Intuitively, the effect of obstacle encounters seems likely to involve some ballistic component – i.e., an animal’s motion is redirected or slowed. However, the observed effect may deviate from intuition based on the animal’s mechanics and behavior, as well as details of the obstacle encounter, such as the initial motion of the animal and which body structures contact the obstacle [13]. Tolerance for collisions may vary between taxa – for instance, birds of prey can suffer bone fractures that eventually heal, whereas wing damage in insects is permanent [1,15]. Because wing damage can increase mortality in insects, the avoidance and consequences of wing collisions has been emphasized in many insect flight studies [9,10,16–18]. Furthermore, numerous insect species have wing morphologies that minimize damage by flexibly deforming during obstacle encounters, and these features have become the focus of studies aimed at extracting wing designs for bio-inspired flying vehicles [19–21]. As a result, our knowledge about obstacle encounters in flying insects is heavily focused on a specific anatomical structure, the wing, even though obstacle encounters and injuries can occur to other parts of the body [1]. We therefore know little about the full range and consequences of obstacle encounters that occur during insect flight.

Here, we use the Valley Carpenter Bee, *Xylocopa varipuncta*, to determine the types of obstacle encounters that can occur in flying insects and their consequences for performance. Among bees (family Apidae), carpenter bees in the genus *Xylocopa* are exceptionally large (wing span > 4 cm) and are important pollinators of crops and wild plants [22]. Thus, they commonly face the challenge of maneuvering a large body through dense foliage. *Xylocopa* spp. are also models for physiological and neurobiological studies due to their large flight muscles and visual acuity [22,23]. We used high-speed video cameras to film *X. varipuncta* flying through narrow gaps in an obstacle course with varying environmental conditions, including moving vs. stationary obstacles and wind vs. still air, in a laboratory flight tunnel. Using these data, we answered the following questions: (1) How frequent are different kinds of obstacle encounters?, What environmental factors affect the likelihood of encounters?, and (3) What are the performance consequences of each kind of encounter?

## Methods

Female carpenter bees (*Xylocopa varipuncta, n* = 15) were collected from the University of California, Davis campus and used immediately in flight experiments. Individual bees were placed in a flight tunnel (20 × 19 × 115 cm; width x height x length) [10,24], which included a series of vertical obstacles that spanned the middle of the tunnel (obstacle diameter = 7 mm, space between obstacles = 34.44 ± 2.80 mm; mean ± SD). The bees’ wing spans (45.19 ± 2.11 mm, tip to tip) were larger than between obstacles; thus, bees needed to rotate their body (e.g., yaw) to pass between obstacles. Obstacles were attached to a mechanical arm that oscillated laterally (amplitude = 21 mm, frequency = 2 Hz) or remained stationary. Fans at each end of the tunnel could be turned on to produce a gentle breeze (mean velocity = 0.54 m/s) or off for still air. Wind direction was constant: bees flying in one direction experienced headwinds and in the other direction tailwinds. Up to 12 flights through the obstacles were elicited per bee, using full spectrum lights at each end of the tunnel [10,24]. Obstacle motion (stationary versus moving) was fixed for a given bee, but all bees experienced wind and still air, with wind condition switched after approximately six flights and the order of wind conditions alternated between bees. Thus, test conditions were still air with stationary obstacles (*n* = 40 flights) or moving obstacles (*n* = 34), and wind with stationary obstacles (*n* = 42) or moving obstacles (*n* = 29).

Flights were filmed at 1500 frames/s with two synchronized Phantom v611 cameras (Vision Research, Inc., Wayne, NJ, USA) positioned 30º from the vertical on opposite sides of the obstacles. Cameras were calibrated using a standard checkerboard calibration method and MATLAB functions [25,26]. In each video, the positions of the bee’s head (midpoint between antennae), thorax (approximating the body centroid), and wing tips were tracked with the machine-learning software DeepLabCut [27]. Tracked points were checked and manually corrected, and obstacle positions labeled using DLTdv6 in MATLAB [28]. Labeled positions were converted from two-dimensional coordinates in each camera view into three-dimensional space using MATLAB functions.

We classified and counted each obstacle encounter. The most common encounters were body collisions (head, thorax, or abdomen contacted obstacles), leg collisions (one or more forelegs contacted obstacles), and wing collisions (the distal half of one or more forewings contacted obstacles) (Fig. 1).

**Figure 1.**
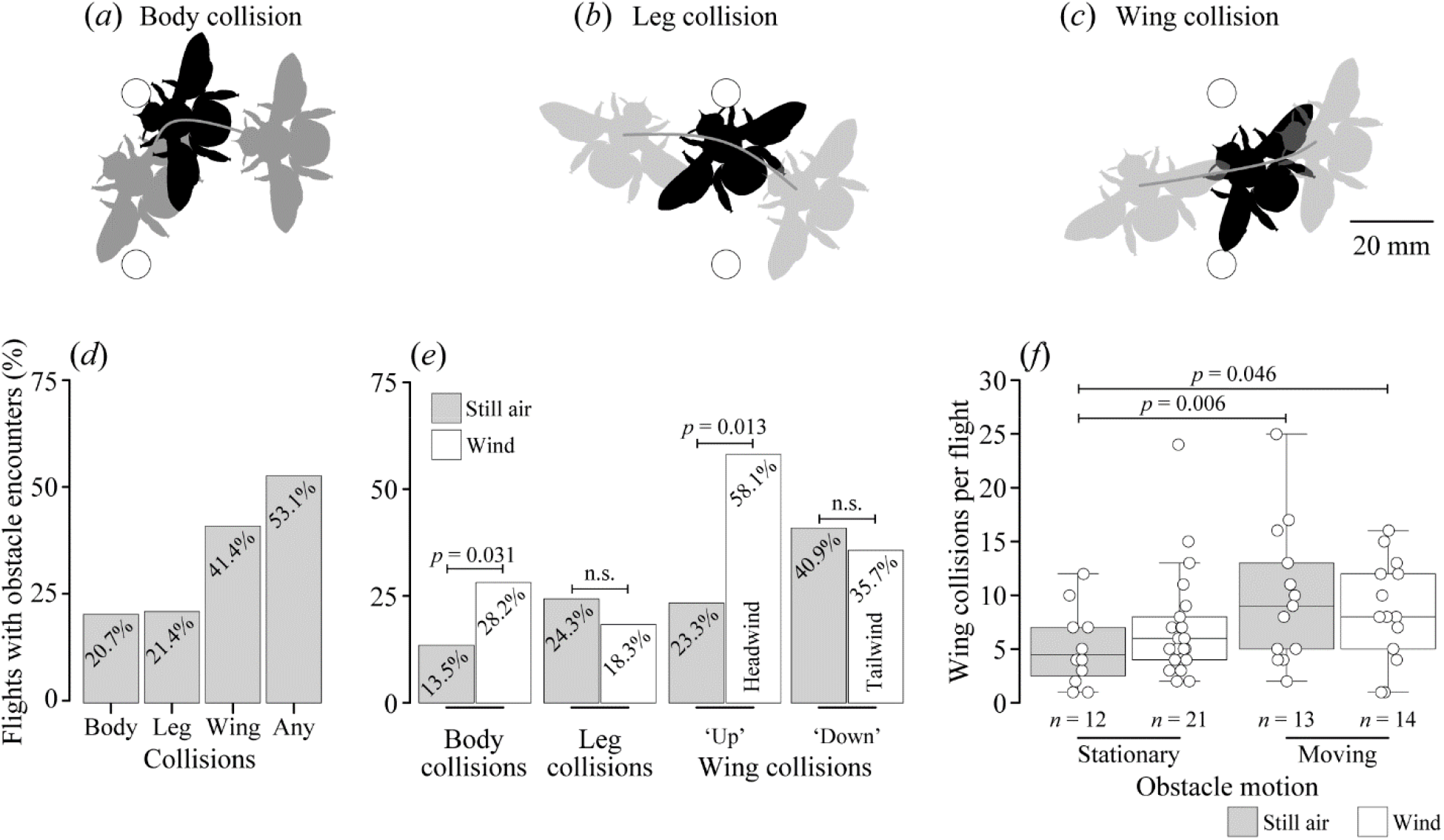
Types of obstacle encounters observed in carpenter bees. Top row: real examples of body, (*b*) leg, and (*c*) wing collisions in bees flying past obstacles from right to left (see supplementary movies). Black outlines show the moment of each encounter. Gray outlines show body positions 20 ms before and after encounters. White circles show obstacle positions. Bottom row: (*d*) frequencies of encounter types, and (*e*) frequencies grouped by wind condition and, for wing collisions, by flight direction. ‘Up’ and ‘Down’ refer to separate directions in the flight tunnel. (*f*) Number of wing collisions per flight (excluding flights without wing collisions), grouped by wind condition and obstacle motion. Brackets show statistical comparisons (*p* < 0.05 for significance; ‘n.s.’ = not significant). Only environmental factors (i.e., wind, flight direction, obstacle motion) retained by the models are shown.

To test which experimental conditions contributed to the occurrence of encounters, we used a generalized linear mixed-effects model (GLMM). Models were implemented as logistic regression models with the ‘glmer’ function in the R package *lme4* [29], with variables for wind (yes versus no), flight direction (upstream versus downstream), and obstacle motion (moving versus stationary). Bee identity was included as a random effect to account for multiple observations per individual. We allowed for statistical interactions between all experimental variables and generated alternative models by removing terms in a stepwise selection process. Models were compared by their Akaike’s Information Criterion (AIC) via the ‘AIC’ function in the R package *stats* [30]. Models with the lowest AIC were evaluated with the ‘Anova’ function from the R package *car* [30].

Among flights with obstacle encounters, there was wider variation in the number of wing collisions per flight (range = 1-25) compared to body collisions (range = 1-2) or leg collisions (maximum = 1). We tested which experimental variables best predicted the number of wing collisions per flight by using a GLMM with a Poisson distribution on flights with at least one wing collision. Model selection and evaluation were carried out as described above. Post-hoc comparisons of model terms were conducted with Tukey HSD tests using the ‘lsmeans’ function in the R package *emmeans* [33].

We assessed how obstacle encounters affected flight performance. In every video, we identified the first occurrence of each encounter type and defined a 20-ms period before and after each event. This temporal window allowed us to quantify performance immediately before and after encounters, as in [20]. Occasionally, pre- and post-encounter periods contained additional collisions, a common outcome when flying near clutter, but the narrow analysis window allowed us to examine changes in flight performance primarily occurring around the focal obstacle encounter. Videos yielded either one, two, or three encounter types (*n* = 42, 26, and 9 flights, respectively).

For each encounter, we measured the change in horizontal ground speed and yaw angle between the pre- and post-encounter periods, as well as the post-encounter yaw rate, where yaw angle was the body angle about the vertical axis. To calculate kinematics, we smoothed three-dimensional position data with cubic smoothing spline curves via the ‘smooth.spline’ function in *stats*. Horizontal ground speed was calculated as the change in x-y position (lateral and longitudinal movements, omitting vertical motion) per time. Yaw was calculated by converting the Cartesian coordinates of the head and thorax to spherical coordinates via the ‘cart2sph’ function in the R package *pracma* [31] and finding the horizontal angle between the body points and the tunnel’s long axis. Yaw rate was calculated as change in yaw per time.

We used a linear mixed-effects model to test whether the change in flight metrics depended on encounter type, wind condition, and/or obstacle motion. Models were implemented with the ‘lme’ function in the R package *nlme* [32]. Model selection, evaluation, and post-hoc comparisons were carried out as described above. Assumptions of normality and homogeneity of variances were checked with Shapiro’s Tests and Levene’s Tests, respectively. When necessary, variance structures of model terms were modified using the ‘varIdent’ function in *nlme*.

## Results

Of the 145 recorded flights from *Xylocopa varipuncta*, 20.7% (*n* = 30) included a body collision, 21.4% (*n* = 31) included a leg collision, and 41.4% (*n* = 60) included a wing collision; overall, more than half of flights (53.1%, *n* = 68) included some type of obstacle encounter (Fig. 1d). Body collisions were more likely to occur in wind (frequency = 28.2%, *n* = 20/71 flights) versus still air (13.5%, *n =* 10/74 flights) (GLMM: χ^2^ = 4.673, df = 1, *p* = 0.031), whereas leg collisions showed similar frequencies between wind (18.3%, *n =*13/71 flights) and still air (24.3%, *n* = 18/74 flights; Fig. 2e) (χ^2^ = 1.426, df = 1, *p* = 0.232). Wing collision frequency depended on wind and flight direction (χ^2^ = 6.341, df = 1, *p* = 0.012), such that flights in headwinds (but not in tailwinds) were more likely to contain wing collisions (58.1%, *n =* 25/43 flights) than flights in the same direction with still air (23.3%, *n* = 7/30 flights; Fig. 2e) (Tukey HSD tests: *p* = 0.013). Notably, our AIC-based model selection process indicated that obstacle motion was not a strong predictor of the likelihood of any encounter type occurring.

**Figure 2.**
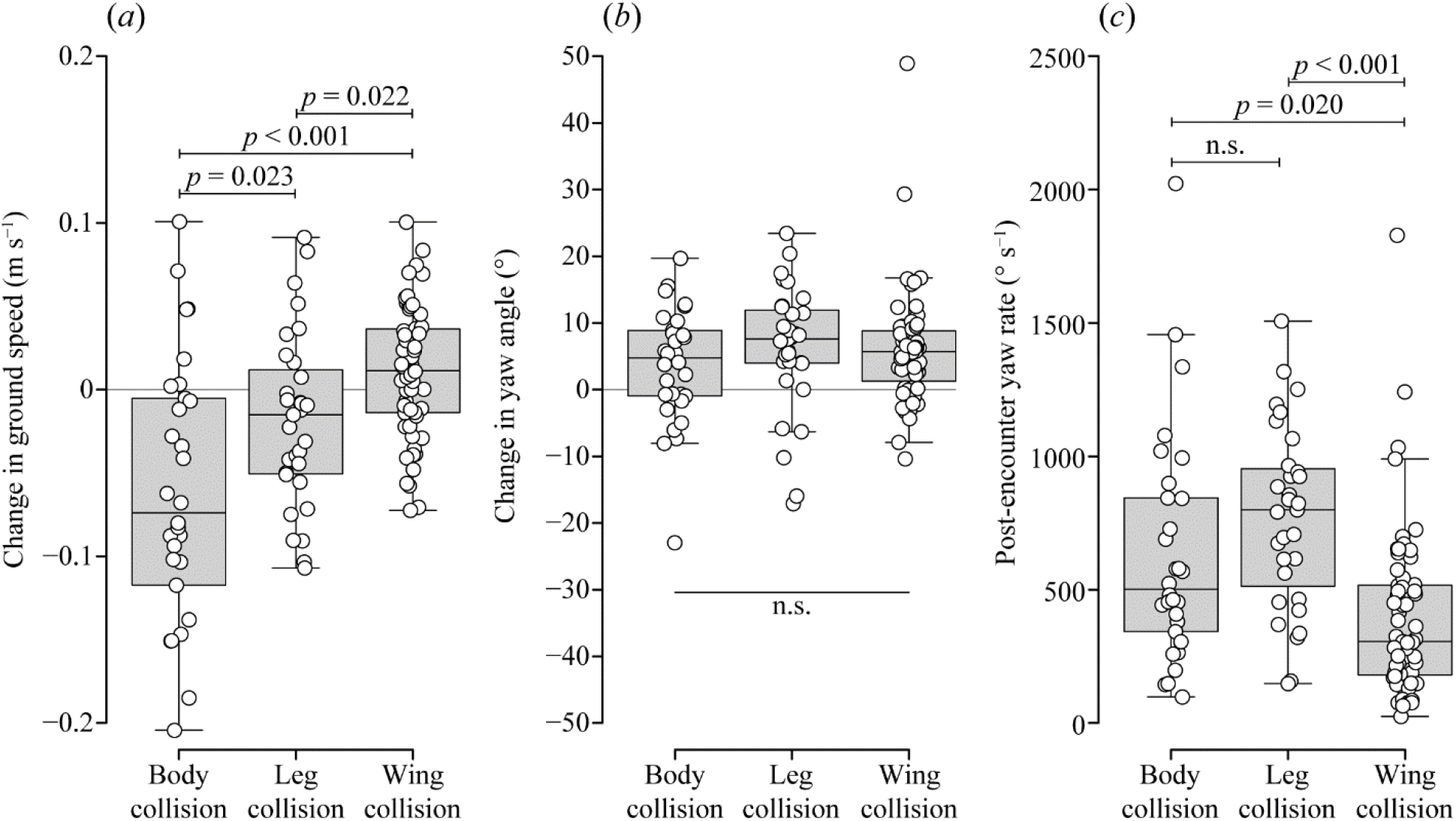
Effects of obstacle encounters on flight. Changes in (*a*) horizontal ground speed and *(b)* yaw angle between the pre- and post-encounter periods, and (*c*) post-encounter yaw rate (*n* = 30 body collisions, 31 leg collisions, 60 wing collisions). Increases in yaw (*b*) indicate rotations away from the tunnel’s centerline. Brackets show statistical comparisons (*p* < 0.05 for significance; ‘n.s.’ = not significant).

Among flights with wing collisions, the number of wing collisions per flight depended on wind and obstacle motion (χ^2^ = 7.011, df = 1, *p* = 0.008; Fig. 1f). Flights in still air with stationary obstacles had fewer wing collisions (5.3 ± 3.5 wing collisions; mean ± SD) than flights in still air with moving obstacles (9.9 ± 6.5 wing collisions) (Tukey HSD test: *p* = 0.006) or flights in wind with moving obstacles (8.6 ± 4.8 wing collisions) (*p* = 0.046).

Body, leg, and wing collisions (Fig. 1a-c) each had distinctive effects on ground speed, with large decreases in speed after body collisions (−0.07 ± 0.08 m/s), small decreases after leg collisions (−0.02 ± 0.05 m/s), and small increases after wing collisions (0.01 ± 0.04 m/s) (Fig. 2a)(χ^2^ = 25.896, df = 2, *p* < 0.005; Tukey HSD tests: *p* < 0.05).

Changes in bees’ yaw angle resulting from collisions were not affected by encounter type (χ^2^ = 0.574, df = 2, *p* = 0.751) (Fig. 2b) but were affected by wind, with larger changes in yaw angle for bees flying in still air (7.29 ± 6.12º) versus wind (3.96 ± 10.33º) (χ^2^ = 7.378, df = 1, *p* = 0.007). Post-encounter yaw rate depended on encounter type (χ^2^ = 21.284, df = 2, *p* < 0.005) (Fig. 2c), with slower rates after wing collisions (305.6 [333.6] º/s; median [interquartile range]) versus body collisions (501.5 [491.4] º/s) (Tukey HSD test: *p* = 0.020) and leg collisions (799.2 [440.3] º/s) (*p* < 0.005) (Fig. 2c).

## Discussion

We found that carpenter bees experienced three distinct types of obstacle encounters when traversing challenging flight environments. Wing collisions were the most frequent encounter (occurring in about 40% of flights), but body and leg collisions each occurred in approximately 20% of flights as well. These results suggest that collisions with obstacles may be common for insects flying through natural, complex environments, such as cluttered vegetation, and that these encounters are diverse – both in terms of the environmental conditions that increase their likelihood and in their effects on flight performance.

Wind increased the likelihood of some encounters, but this effect was not uniform. Body collisions occurred more frequently in wind, but leg collisions were not affected – and wing collisions occurred more frequently only for bees flying into a headwind. Obstacle motion did not affect the likelihood of any type of encounter, but obstacle motion (in still air or in wind) did lead to a greater number of wing collisions for flights in which wing collisions occurred. Given that our experiment tested only one obstacle arrangement, one type of obstacle motion, and one mild wind speed, it is clear that additional studies would greatly improve our understanding of how environmental variables affect the likelihood and consequences of obstacle encounters.

Most insect flight studies focus on collision-free flight, yet in our study more than half of the flights recorded (53.1%) contained at least one type of collision, suggesting that additional research is needed on flights involving obstacle encounters. In addition, most previous studies addressing collisions have focused only on wing collisions, in part because wing collisions lead to cumulative damage that can impair flight performance over time. However, our work suggests that other types of obstacle encounters, such as body and leg collisions, may have more important immediate effects on flight performance, leading to larger reductions in flight speed and larger changes in body rotation rates than wing collisions. Overall, our work suggests that the incidence, diversity, and potential effects of obstacle encounters experienced by flying insects remains vastly underexplored in the scientific literature.

## Supporting information

Example video of a body collision

Example video of a leg collision

Example video of a wing collision

## Data accessibility

Data are available from the Dryad Digital Repository https://doi.org/10.25338/B8M939.

## Authors’ contributions

**Nicholas P. Burnett:** Conceptualization, Methodology, Investigation, Formal analysis, Data curation, Writing – original draft, Visualization, Funding acquisition. **Stacey A. Combes:** Conceptualization, Methodology, Supervision, Project administration, Writing – review & editing.

## Acknowledgements

The authors thank M. Badger for assistance with DeepLabCut and MATLAB.

